# KoT: an automatic implementation of the *K/θ* method for species delimitation

**DOI:** 10.1101/2021.08.17.454531

**Authors:** Yann Spöri, Fabio Stoch, Simon Dellicour, C. William Birky, Jean-François Flot

**Affiliations:** Evolutionary Biology & Ecology, Université libre de Bruxelles (ULB), Brussels, Belgium; Interuniversity Institute of Bioinformatics in Brussels – (IB)^2^, Brussels, Belgium; Spatial Epidemiology Lab (SpELL), Université libre de Bruxelles (ULB), Brussels, Belgium; Department of Microbiology, Immunology and Transplantation, Rega Institute, Laboratory for Clinical and Epidemiological Virology, KU Leuven, University of Leuven, Leuven, Belgium; Department of Ecology and Evolutionary Biology, The University of Arizona, Tucson, Arizona, USA

**Keywords:** 4X rule, *K/θ*, species delimitation, molecular systematics, DNA taxonomy

## Abstract

*K/θ* is a method to delineate species that rests on the calculation of the ratio between the average distance *K* separating two putative species-level clades and the genetic diversity *θ* of these clades. Although this method is explicitly rooted in population genetic theory, it was never benchmarked due to the absence of a program allowing automated analyses. For the same reason, its application by hand was limited to small datasets of a few tens of sequences.

We present an automatic implementation of the *K/θ* method, dubbed KoT (short for “K over Theta”), that takes as input a FASTA file, builds a neighbour-joining tree, and returns putative species boundaries based on a user-specified *K/θ* threshold. This automatic implementation avoids errors and makes it possible to apply the method to datasets comprising many sequences, as well as to test easily the impact of choosing different *K/θ* threshold ratios. KoT is implemented in Haxe, with a javascript webserver interface freely available at https://eeg-ebe.github.io/KoT/

## Introduction

Methods to delineate species from sets of DNA sequences have been an intense field of research for the last 20 years (Sites and Marshall, 2003; Flot, 2015). Some methods delimit species based on phylogenetic trees, others on genetic distances, and yet others on allele sharing (Fontaneto et al., 2015). Among these methods, one called *K/θ* (Birky et al., 2010; Birky, 2013; Birky and Maughan, 2020) stands out by resting on the genealogical species concept, in which closely related populations are considered as distinct species when their lineages for a given locus (or set of loci) are reciprocally monophyletic, that is, “if their loci coalesce more recently within the group than between any member of the group and any organisms outside the group” (Baum and Shaw, 1995). Of course, sampling all lineages from a population is usually impossible, but population genetic theory provides ways to calculate the probability that the lineages of two populations are monophyletic given the observation that the sequences sampled from these populations form clades in a phylogenetic tree (Hudson and Coyne, 2002).

In particular, Rosenberg (2003) provides a formula that uses Watterson’s estimator of genetic diversity *θ* = 4*N_e_μ* (Watterson, 1975) of each of two clades of sequences, the number of sequences in each of them, and the mean pairwise sequence difference *K* between the two clades to calculate the probability that the corresponding two populations are reciprocally monophyletic, i.e. distinct species according to the genealogical species concept. This formula is complex, but when the two *θ* values are similar and the number of sequences in each clade is higher than three, a useful rule of thumb is that pairs of clades with a *K/θ* ratio higher than 4 have a probability of at least 0.95 of belonging to different species. This forms the basis of the so-called “4X rule” (Birky and Barraclough, 2009), which has been widely used to delineate species in a variety of organisms. However, one may wish to choose a more stringent threshold: for instance, a *K/θ* ratio higher than 6 entails a probability of monophyly higher than 0.99 (according to equation 9 in Rosenberg, 2003).

Despite the theoretical appeal of this method based on an explicit criterion inspired by population genetic, its practical application has been hampered by the lack of a program performing the needed calculations automatically. To fill this gap, we introduce here KoT (short for “K over Theta”), an automatic implementation of the *K/θ* method using the programming language Haxe (Dasnois, 2011).

## Description

KoT takes as input an alignment of DNA sequences in the FASTA file format. Users can specify the *K/θ* threshold they wish to use to delineate species (by default, 4).

KoT starts by calculating all the pairwise nucleotide distances among the sequences in the dataset. In case the input contains indels and/or missing data (encoded respectively as “-” and as “N” or “?”), users can either ask KoT to filter out completely the corresponding positions in the alignment (“complete deletion” mode) or to retain them (“pairwise deletion” mode, in which case positions with missing or ambiguous data are ignored during pairwise comparisons). From this set of pairwise distances, KoT then computes a neighbor-joining (NJ) tree and the *K/θ* ratios of each pair of sister clades using the procedure outlined in Birky and Maughan (2020).

To compute the genetic diversity *θ* of each clade, KoT starts by calculating the nucleotide diversity *π* (Nei and Li, 1979) as the mean of all nucleotide-level differences *π_ij_* (number of nucleotide differences per nucleotide site between sequences *i* and *j*) among the 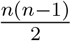 pairs of sequences in a clade of *n* sequences, i.e. 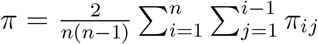 (Equation 10.6 in Nei, 1987). An equivalent way to calculate *π* found in the literature is to compare all pairs of haplotypes instead of all pairs of sequences: in that case, the formula above becomes 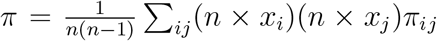 (where *x_i_* is the frequency of haplotype *i* in the clade), which simplifies into 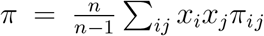 (Equation 10.6 in Nei, 1987), i.e. the average heterozygosity multiplied by a sample size correction *n*/(*n* − 1) (Nei and Tajima, 1981; Korunes and Samuk, 2021). As KoT uses the direct formula comparing pairs of sequences, however, no sample size correction is needed.

For clades made up of identical sequences (in which case *π* = 0), an upper bound of *π* is sought by assuming that one sequence differs from the others by a single mutation, i.e. by replacing one of the 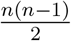 pairwise distances with 1/*L*, where *L* is the sequence length; in such case *π* become 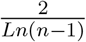 (Birky et al., 2010). As this ratio is not defined for *n* =1, KoT uses *n* = 2 (i.e. *π* = 1/*L*) for clades comprising a single sequence. To estimate the genetic diversity *θ* associated with a specific clade, KoT uses the formula 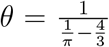 (Equation 9 in Tajima, 1996), which corrects for multiple hits based on the Jukes-Cantor model of sequence evolution (Jukes and Cantor, 1969).

To compute the genetic divergence *K* between sister clades A and B, KoT computes the mean pairwise nucleotide distance 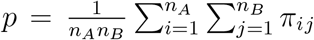 between the sequences in the two clades, where *n_A_* stands for the number of individuals in clade A and *n_B_* stands for the number of individuals in clade B, then corrects it for multiple substitutions using the Jukes-Cantor formula 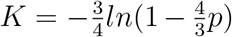 (Jukes and Cantor, 1969). Using the Jukes-Cantor correction for calculating *K* is important to ensure that both terms of the ratio *K/θ* are computed using the same evolutionary model.

Finally, KoT computes the *K/θ* ratios of clades A and B and compares the smallest of the two with the threshold chosen by the user to delineate species. These calculations are performed iteratively from the leaves of the tree all the way to its root. When a ladder structure ((A,B)C) or a polytomy (A,B,C) is encountered, KoT starts by comparing the two clades separated by the smallest distance *K*, i.e. A and B: if the result of the calculation does not support the hypothesis that A and B are different species, the A+B clade is then compared to C to find out whether they are conspecific or heterospecific; whereas if the result of the calculation indicates that A and B are likely two distinct species, C is compared with whichever of A and B has the smaller average distance *K* to C (Birky et al., 2010). If C is then deemed to be distinct as well, the final result returned is three species A, B and C. On the other hand, if the result of the calculation suggests that C is conspecific with e.g. B, an additional comparison of C with A is warranted to ensure the transitivity of the result (Dellicour and Flot, 2018). As the extra calculations this entails can take lots of time in complex cases, this comparison is only performed if the box “transitivity” is checked by the user prior to running the analysis. If the “transitivity” box is not checked (as by default), KoT simply returns in such cases two species A and B+C; when the “transitivity” box is checked, by contrast, KoT checks whether the *K/θ* ratio for the C vs. A comparison is also above the user-selected threshold: if so, the final delimitation returned is a pair of species A and B+C; whereas if the calculation does not support the monophyly of A vs. C, the final result returned is a single species A+B+C.

KoT outputs a tree in which the *θ* values of each pair of clades being compared are displayed on the tree next to the node uniting them, together with their *K* distance and the *K/θ* ratio (obtained using the larger of the two *θ* values). Colors are applied to the tree in order to visually delineate the different species. A partition list, i.e. a two-column table indicating, for each sequence in the input FASTA, the species to which it was attributed (Spöri and Flot, 2020), is also outputted below the tree where is can be easily copied/pasted into other applications.

## Biological Example

To investigate the behavior of KoT, we reanalyzed the COI dataset from one recent article (Stoch et al., 2020). In this article, a dataset of 34 COI sequences of specimens of the *Niphargus tatrensis* species complex was analyzed using a diversity of approaches: mPTP Kapli et al. (2017) delimited seven putative species, ABGD (Puillandre et al., 2012) returned ten of them, and bPTP (Zhang et al., 2013) and ST-GMYC (Pons et al., 2006) delineated eleven species-level units. The methods chiefly differed in their delimitation of species among the non-Austrian specimens included in the study but were largely congruent in their treatment of the Austrian specimens, with mPTP, bPTP and ST-GMYC finding four species and ABGD delimiting five species in Austria.

When run with a *K/θ* threshold ratio of 4 (Figure 1), KoT returned twelve species, including five for Austria (separated by *K/θ* ratios of 21.39, 17.50, 4.52 and 5.98); with a *K/θ* threshold ratio of 5 (Figure 2), the method returned eleven species, with precisely the same putative boundaries as those obtained using bPTP and ST-GMYC (including four species for Austria separated by *K/θ* ratios of 21.35, 17.44 and 5.94); finally, with a *K/θ* threshold ratio of 6 (Figure 3) KoT returned eight species-level units, notably lumping together all Austrian specimens into a single putative species. This highlights the sensitivity of this method to the *K/θ* threshold parameter.

**Fig. 1.**
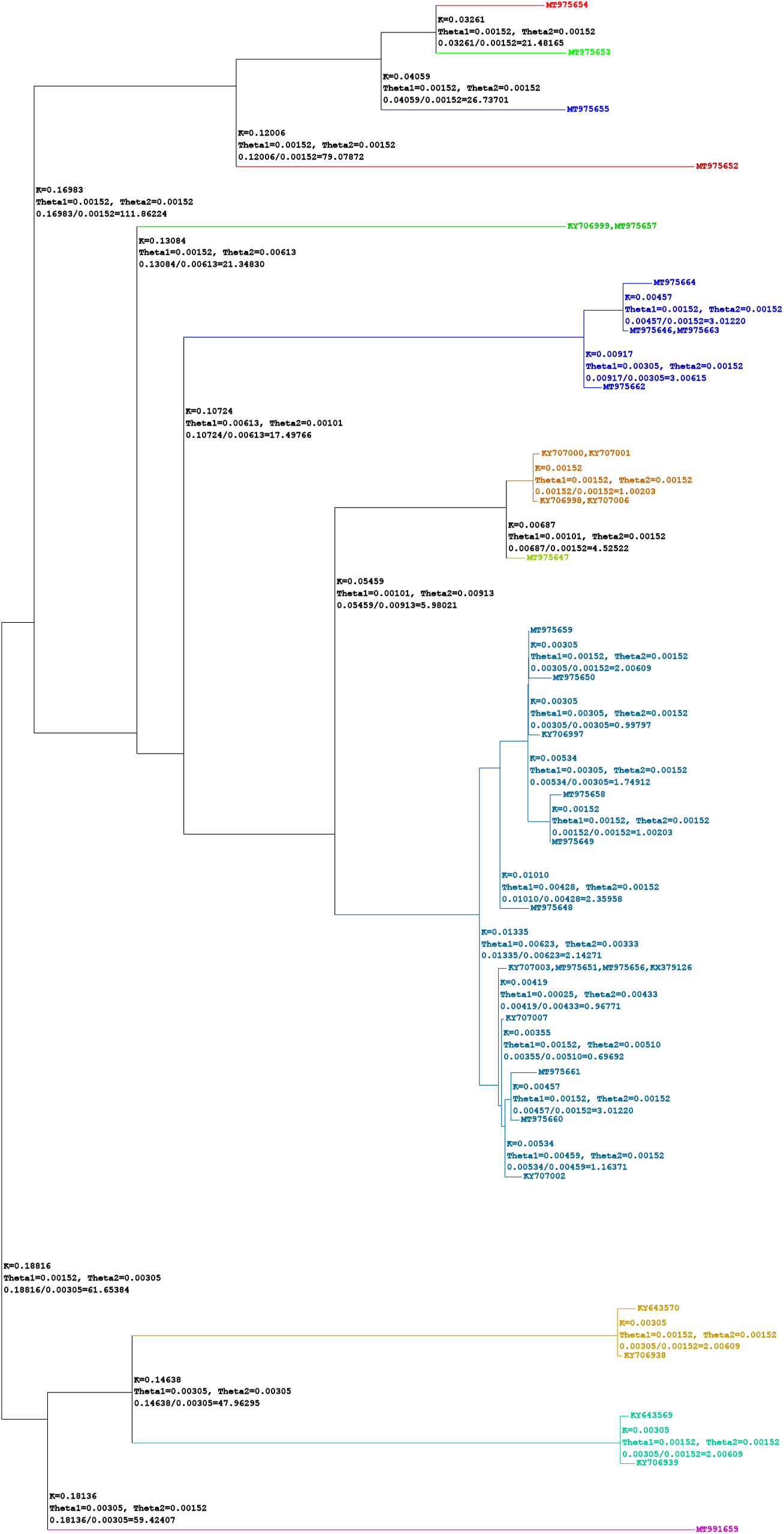
Output of KoT when run on the COI dataset of Stoch et al. (2020) with a *K/θ* threshold of 4

**Fig. 2.**
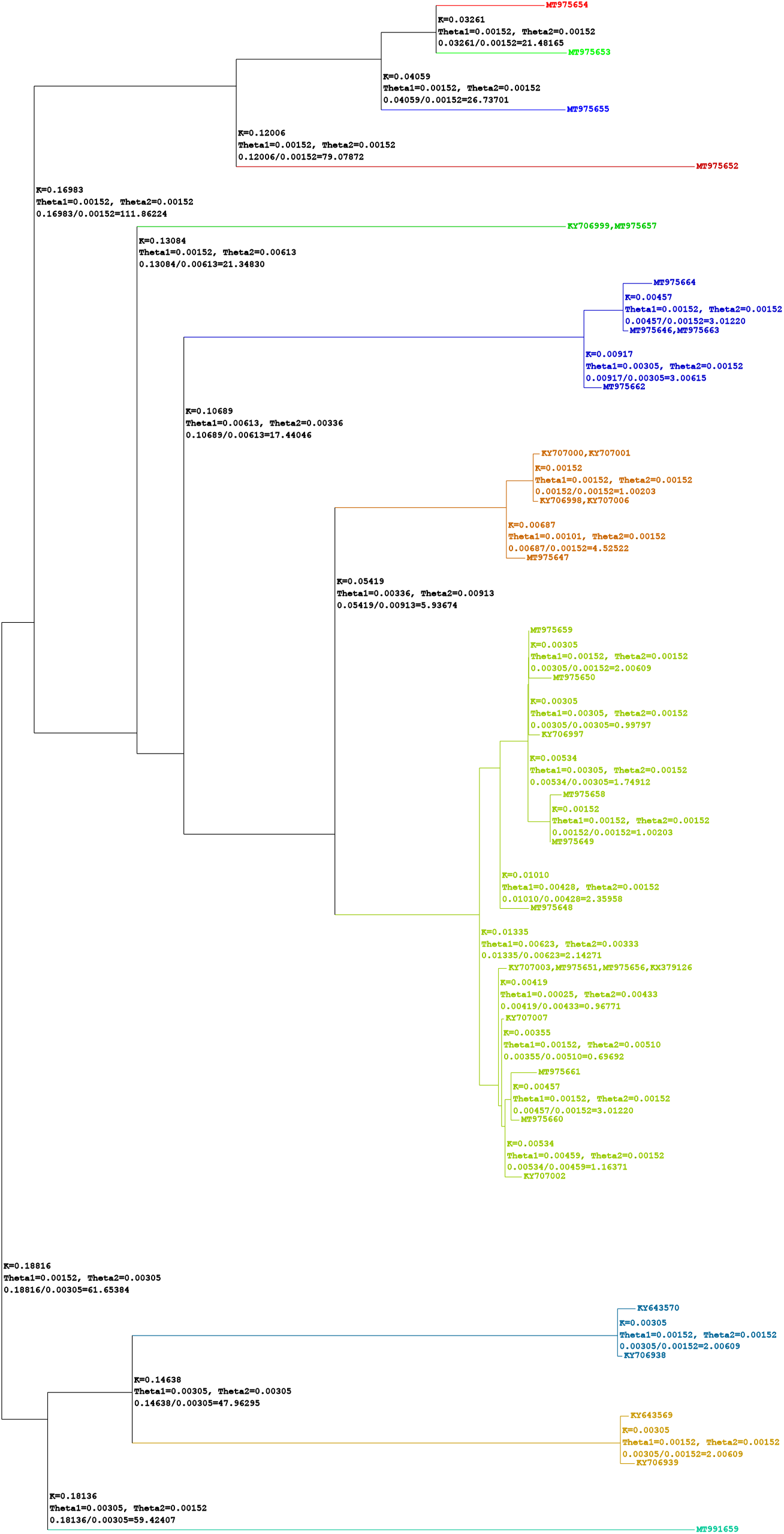
Output of KoT when run on the COI dataset of Stoch et al. (2020) with a *K/θ* threshold of 5

**Fig. 3.**
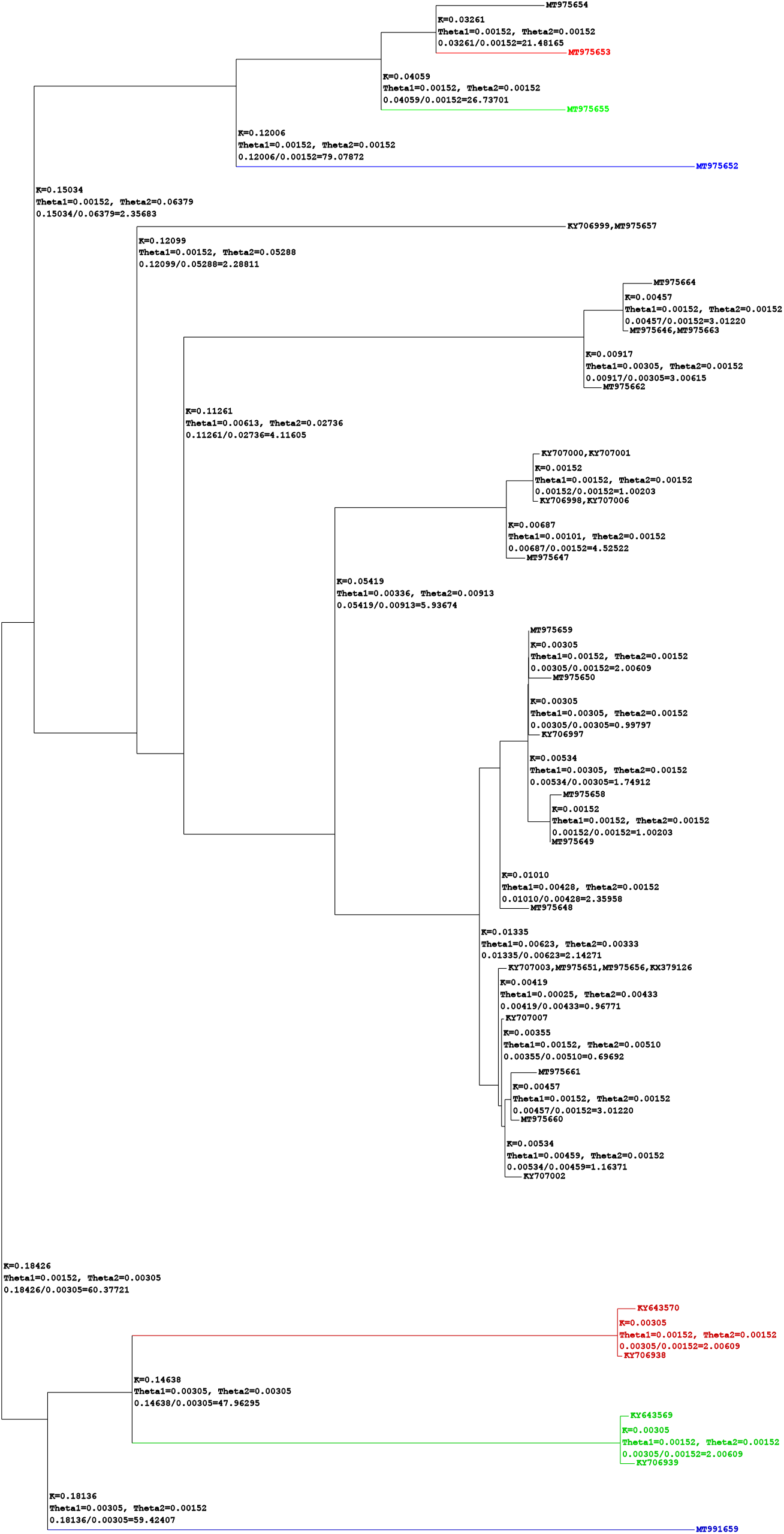
Output of KoT when run on the COI dataset of Stoch et al. (2020) with a *K/θ* threshold of 6

## Availability

KoT is written in Haxe. Its source code is available at https://github.com/eeg-ebe/KoT, and a javascript webserver is freely accessible at https://eeg-ebe.github.io/KoT/.

## Notes

### Competing Interest Statement

The authors have declared no competing interest.

### Summary of Updates

Typos corrected (including an important one in the formula for the calculation of K - this typo was only in the paper, the calculations performed by the program were not affected).

